# An Antioxidant Polysaccharide from *Ganoderma lucidum* Induces Apoptotic Activity in Breast Cancer Cell Line

**DOI:** 10.1101/2022.01.04.474971

**Authors:** Md. Moyen Uddin Pk, Rumana Pervin, Mohammad Shahangir Biswas, Md. Matiar Rahman

## Abstract

The purpose of this study is to elucidate the apoptotic activity of *Ganoderma lucidum* polysaccharide (GLP) in a human breast cancer cell line MCF-7 *in vitro*. According to DPPH assay, GLP showed a good antioxidant (IC_50_ value is 202.4 μg/mL). Based on MTT assay, the results showed that GLP inhibits MCF-7 cells proliferation in a dose- and time-dependent manner (p<0.001). IC_50_ values of the cytotoxicity of GLP and doxorubin were 110.907 μg/mL and 58.206 μg/mL respectively. The results from the flow cytometry indicated that GLP could induce apoptotic activity through inducing the up-regulation of the Bax and Caspase-9 and the down-regulation of the BcL-2 in MCF-7 cells. At 2×IC_50_, GLP increased the early-apoptotic and dead cells of MCF-7 from 18.23% to 34.76% and 8.45% to16.34% respectively. In conclusion, the GLP shows anticancer activity against MCF-7 through preventing the proliferation and inducing the apoptosis of MCF-7 cells. Our data provide the potential molecular targets in cancer prevention and reveal the key barriers in the current anticancer drugs development.

## Introduction

Breast cancer is the leading cause for women’s mortality in the world(1). Nearly one quarter of female cancer patients are diagnosed with breast cancer(2). When breast cancer progresses, cancer cells release antiapoptotic factors. Breast cancer continues to be a major concern and threat to human health (3). The treatment options are limited by various subtypes of breast cancer, each requiring different therapy regimens. Chemotherapy and radiotherapy are the standard treatment for breast cancer; however, the treatment results are mostly disappointing for patients. In this situation it is important to explore various alternative therapies for the treatment of breast cancer. In many cultures it is popular to use herbs to treat malignancies, as there are the plenty of anti-cancer compounds in some plants and are used in the manufacture of various modern drugs(4). Different sources of polysaccharides for their anti-cancer activities have been identified(5, 6). Through its possible biological functions, antioxidant(7), anti-tumor(8) and hepatoprotective effects(9), polysaccharides have attracted a lot of attention. Several studies showed that biologically active compounds isolated from mushrooms are primarily linked to polysaccharides for the anti-cancer properties(10–14). Previous researches have shown that polysaccharides isolated from the *Ganoderma lucidum* show antitumor properties(15). As far as we know, no research on the anti-tumor activities of *Ganoderma lucidum* polysaccharide on MCF-7 cells was conducted in Bangladesh. This study aimed to investigate the antitumor effects of *Ganoderma lucidum* polysaccharide (GLP) on MCF-7 cells *in vitro* models. This study was intended as a scientific basis for work on polysaccharide and for the application in medicine. The data support the potential use of GLP for the treatment of breast cancer.

## MATERIALS AND METHODS

### Chemicals

All culture media including RPMI-1640, fungizone, FBS, and penicillin-streptomycin solution, DEAE, DPPH, ABTS, MTT, Tris, DTT, SDS, DCFH-DA, ethanol, acetone, and sulfuric acid were obtained from Sigma-Aldrich, USA. All other chemicals including KBr, butanol, chloroform, phenol, EDTA, trypsin, ether, glycerol, hydrogen peroxide were purchased from Merck, Germany. In addition, FITC/PI, Bcl-2, Bax, Cas-9 and colorimetric assay kits were obtained from Abnova, Germany. Western blot antibodies against Bax, Bcl-2 and Caspase-9 were purchased from Abnova (Germany), CST (Boston, MA), SCB (Stanta cruz, CA) respectively.

### Sample collection

Fruiting bodies of *Ganoderma lucidum* were collected from the Mushroom Development Institute, Bangladesh. It was authenticated by Scientific officer at National Mushroom Development and Extension Center, Savar, Dhaka-1340 in Bangladesh. No.: SR-0022(2/3/2018)

### Extraction

Polysaccharide from the *Ganoderma lucidum* (GL) fruiting bodies was extracted according to the methods of our previously published article^9^. In short, the dried powder (30 g) of the *Ganoderma lucidum* (PO) was extracted with distilled water (1:30) at 90°C for 3 h under stirring to remove hot water dissolved pigments, polyphenols, and monosaccharides. The liquid fraction was collected by centrifugation at 5000 rpm for 15 min at 56°C and precipitated with 95% ethanol (1:3, v/v) for 12 h at 4°C. Precipitates were observed and collected by centrifugation (4500 rpm for 20 min at 4°C) and washed consecutively with acetone (1:3 v/v, 95%), ether (1:3 v/v, 95%), and ethanol (1:3 v/v, 95%) at 4°C. The precipitate was dried in vacuum to obtain protein-bound Polysaccharide (PBP). The total protein content and the total polysaccharides content were quantified by the Lowry method (16) and phenol-sulfuric acid method(17) respectively.

### Protein free polysaccharide isolation

GLP crude extract was deproteinated using Sevag method(18). According to Sevag principle, the sample and Sevag reagent ratio was 2:1(Chloroform:n-Butanol, 5:1). The optimized ratio of GLP to Sevag reagent was 2:1.

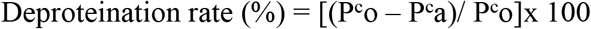

where P^c^o; The content of protein before deproteinization, P^c^a; The content of protein after deproteinization

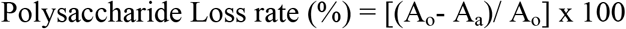

where A_o_; polysaccharide content before deproteinization, A_a_; polysaccharide content after deproteinization

After deproteinization, the solution was desalinated with hyperfiltration (3 Kda, 5.0 mL/min for 8 h) and the filtrate was mixed with 99.5% ethanol and the mixture was kept at 4°C for 24 h.

After centrifugation (4000 rpm, 4°C for 15 min), the precipitate was dissolved in dH_2_O and then lyophilized in a vacuum freeze dryer to obtain the crude polysaccharide, GLP. The standard curve was constructed using D-glucose (C_6_H_12_O_6_) and the linearity of standard curve and extraction yield of GLP were expressed as follows:

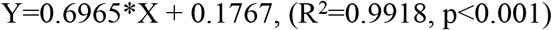

where Y represents the absorbance; X represents the GLP content (mg/mL).

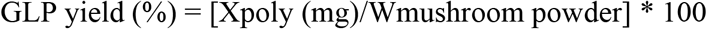

where X represents the GLP content (mg); W represents the weight of the mushroom powder (g).

### Purification of polysaccharide by chromatographic methods

DEAE-Ion exchange chromatography and gel filtration chromatography (GFC) were used to remove fine particles from crude GLP.The crude polysaccharide was loaded onto DEAE-52 column (2.6X80 cm) and the stage gradient elution was carried out using dH_2_O, 0.2 M NaCl, and 0.5 M NaCl respectively. The elution flow rate was 0.8 mL/min and carbohydrates content was checked in each 5 mL test tubes using the phenol-sulfuric acid supplementary method. The largest polysaccharide content fractions were combined, collected and loaded onto Sephadex S-300 column (1.6 x 100 cm) and equilibrated with 0.05 M NaCl. The elution flow rate was 24 mL/hr. The largest fraction was collected and desalted against ddH2O over 5 days using dialysis membrane having a 14000 Da MWCO (Thermo Fisher Scientific). After dialysis, the polysaccharide solution was concentrated and lyophilized to obtain refined polysaccharide.

### Relative molecular weight determination using GFC

Molecular weights of the GLP fraction was determined using Sepharose CL-6B in comparision to the molecular weights of known dextran series (10-2000 kDa, 10 mg/mL). The standard curve was used to calculate Mw of GLP and this curve was constructed at Mw vs K_av_. The K_av_ was calculated using the following formula:

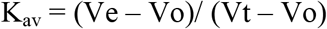

where Vt; total column volume, Ve; elution volume, Vo; void volume, and K_av_; proportion of pores available to the molecule.

The linearity of the standard curve was evaluated with R^2^ value. R^2^ value explains the significant correlation between dependent (K_av_)and independent variables (Mw of dextrans).

### Spectroscopic analysis of polysaccharide

#### UV-vis spectroscopic analysis

GLP was dissolved in distilled water (upto 5%) and were analyzed at 200–600 nm using spectrophotometer (Double beam UV-visible light spectrophotometer, China). In this analysis, the protein and carbohydreate contents were determined.

#### Fourier transform infrared analysis

The fourier transform infrared (FT-IR) method was used to determine the organic functional group of polysaccharide. Before grinding with KBr powder, GLP (3 mg) was dried at 40-45°C and then pressed into a pellet (1 mm) for spectrometer measurement. The wavelength was set by a resolution of 8 cm^−1^ from 4000 to 400 cm^−1^ at 25°C.

### Monosaccharide composition

In this experiment, polysaccharide was hydrolyzed with 2M TFA (2 mL) at 120°C for 3 h. After removing excess TFA, hydrolysate was collected and dissolved in ultrapure water (4 mL) and the mixture was reduced with NaBH_4_ (40 mg) at 25°C. After 3 h, the mixture was neutralized with acetic acid (25 %), and was evaporated the solution with rotary evaporator. At 100°C, acetic anhydrate (3 mL) and pyridine (3 mL) were added in each sealed reaction tube and was kept for 1 h. Standard monosaccharides (D-Mannose, D-Glucose and D-Galactose) were prepared in the same procedure. The solutions were analyzed with an Agilent 6890 GC (Column HP-5, 30 m x 0.32 mm).

### Cell lines

The MCF-7 and MCF-10a cells were obtained from ATCC (American Type Cell Culture Collection), Manassas, USA. The MCF-7 cells were cultured in RPMI-1640 (Sigma, USA) while MCF-10a cells were maintained in Ham’s F-12 (Sigma, USA) and Dulbecco’s modifed eagle medium supplemented with EGF (20 ng/mL), Insulin (10 μg/mL) and hydrocortisone (250 ng/mL). All media were added with 0.5% Fungizone, 10% FBS and 2% penicillin and streptomycin (100 ng/mL).The cells were incubated at 37°C in 90% humidifed incubator with 5% CO_2_ and, after reaching 80% confluence, cells were harvested using 0.25% trypsin-EDTA. The stock solution was prepared in 0.1% DMSO and processed at −20 ° C before use.

#### DPPH radical-scavenging assay

The DPPH^+^ radicals scavenging assay described by Nithianantham *et al* (19) and Uddin Pk *et al* (10)was used to determine the antioxidant activity of GLP. In short, 50 μL samples (31.25 to 1000 μg/mL) were added in reaction tube and following DPPH (0.004%, w/v) was added in each reaction tube. After vortex properly, the mixtures were incubated at 25°C for 30 min in dark place. In this experiment, the blank, the DPPH control, and the positive control were 80 % ethanol, 0.004% DPPH, and vitamin C respectively. The discoloration of reaction in each tube was measured at 517 nm and measurements were taken in three times. DPPH radicals scavenging by extracts and vitamin C was determined using the following formula:

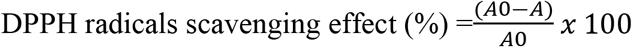

where A0 represents the absorbance of negative control and A represents the absorbance of samples. Results of vitamins C and GLP were reported as EC_50_ using linear regression analysis.

In the EPR analysis, the reaction mixtures were loaded to capillary tubes for the EPR spectrometry (E500, Bruker, USA) and the EPR signals were recorded (60s) with the following EPR measurement conditions: central field 3505±50 g, modulation frequency 100 kHz, modulation amplitude 2.0 g, microwave power 1.02 mW, gain 65 dB, scan time 20.97 s and time constant 40.96 ms

#### Cytotoxicity assay

GLP and doxorubicin (Sigma, USA) were dissolved in DMSO (Sigma, USA) to make a stock solution (1 mg/mL) and stored in −20 °C before use. The working solution was freshly prepared in the complete culture medium. In cytotoxicity study, the MCF-7 cells was seeded at 5 × 10^3^cells/ well (200 μl) into a 96-well plate overnight in 95% humiditiy and 5% CO_2_. After 24 h the cells were incubated with GLP(10 to 1000 μg/mL), negative control, positive control Doxorubin was used as positive control (48 h of incubation). The cells were rinsed with 1× PBS. And then, 20 μl MTT (5 mg/ ml) was added to each well and mixed properly. After 2 h incubation, the dark blue formazan crystals were separated and 100 μl of DMSO was added to dissolve crystals at 37°C for 30 min. All measurements were taken in three times.

Finally, the absorbance was measured at 570 nm using microplate reader (Molecular Devices, USA) and the cell viability (%) was calculated using the following equation:

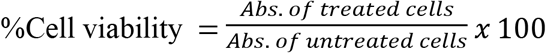

### Study of cell apoptosis

GLP-induced apoptosis was performed by FACS calibur flow cytometer (BD Bioscience, USA). MCF-7 cells (1×10^5^) were seeded in well culture plates and kept overnight. Following day the medium was replaced with new culture medium and cells were treated with GLP (2×IC_50_,) for 48 h. The treated cells were harvested and washed with PBS twice. The cell pellets were re-suspended in binding buffer (500 μL) and stained with Annexin V-FITC and PI using a Dead Cell Apoptosis Kit (Invitrogen) according to the manufacturer’s instruction. After 20 minute incubation at 25°C in the dark, cells were analyzed.

### Cell cycle analysis

Cell cycle phase distribution of MCF-7 cell was evaluated by measuring the DNA content. In this experiment, the treated and untreated cells were trypsinised and washed with PBS. At 4°C, cells were fixed in 70% ice cold ethanol for overnight and washed with PBS to remove residual ethanol. Cells were re-suspended in 300 μl of PBS containing propidium iodide (50 μg/ml), 0.5% Triton X-100 and RNAse A (50 μg/ml) and incubated in water bath at 37 °C for 30 min. Cells were analyzed for cell cycle distribution at different phases (G0/G1, S, G2/M and sub-G1) in FACS Calibur flow cytometer (BD Biosciences, USA).

### Analysis of the mitochondrial membrane potential using the cationic JC-1 dye

In this experiment, previously cultured MCF-7 cells were treated with GLP and incubated for 48 h. Then, cells were treated with JC-1 (20 μl of 200 μl JC-1) for 15 minutes at 37°C. The cells were centrifuged for 5 min at 25°C at 400 x *g*. The cell pellets were washed and re-suspended with PBS. The green JC-1 monomers in mitochondria of apoptic MCF-7 cells was detected in the FL1 channel (FITC, GFP), while the red aggregates of JC-1 in non-apoptotic cells were detected in the FL2 channel (PE, R-PE, RD1).

### Apoptotic protein expression analysis

Apoptotic proteins expression were analyzed using Western blot(11). In this assay, cells were treated with GLP for 48 h and harvested and lysed with assay buffer (Tris 62.5 mM, pH 6.8, DTT 50 mM, 10%SDS, glycerol; Thermo Fisher Scientific, USA). Proteins were extracted and quantified using Bradford assay and separated by 15% SDS-PAGE and transferred onto a polyvinylidene fluoride membrane at 4°C (100 V). After 2 h, the membrane was probed with primary antibody (1:1000) and followed by secondary antibody (HRP; 1:8000). Protein bands were detected by Enhanced chemiluminescence detection kit (Thermo Fisher Scientific, USA).

### Statistical analysis

GraphPad Prism was used for statistical analysis. The data are shown as the mean ± s.d. In order to perform statistical comparisons one-way analysis of variance was used. Effective doses were expressed as IC_50_ values of polysaccharide. A statistically significant difference was known to suggest P<0.05 between the values.

## RESULTS

### Polysaccharide purification and molecular weight determination

Polysaccharide extraction from *Ganoderma lucidum* was carried out using hot-water extraction technique. The validation of the phenolic-sulfuric acid method was carried out using standard D-glucose and the maximum absorbance of D-glucose was recoreded at 490 nm and the linearity of standard curve was 99.18% (p<0.001) (Fig. 1).

**Figure 1.**
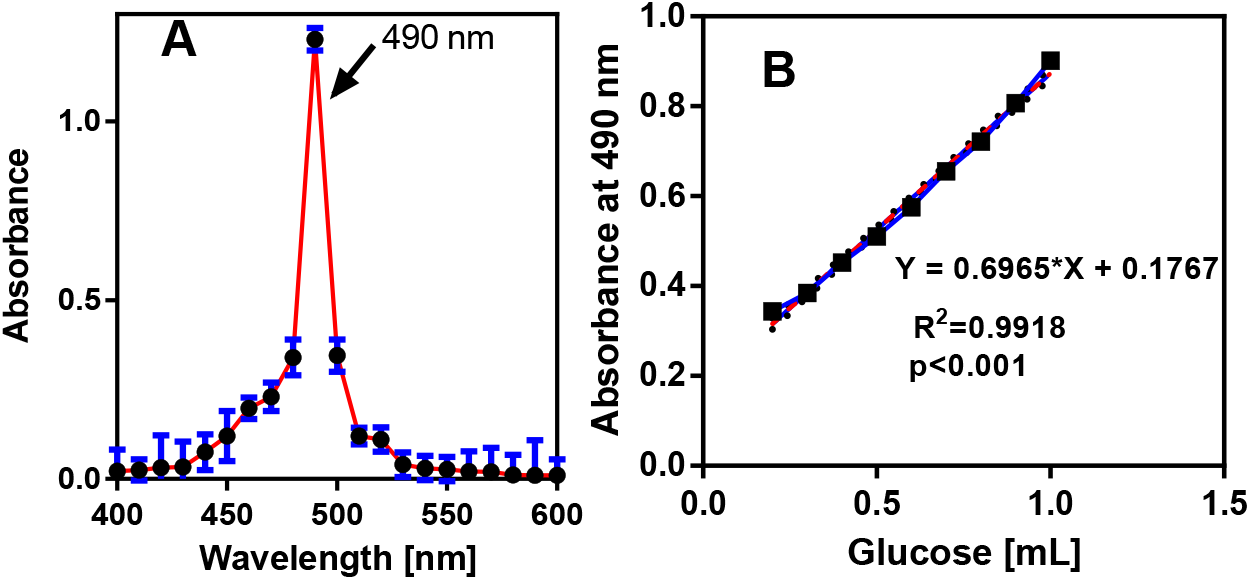
**(A)** Absorbance validation of D-Glucose (range: 400-600 nm) **(B)** The standard curve of D-glucose (Abs. at 490 nm). Y axis shows the absorbance and X axis shows the glucose concentration (mg/mL).

The yield (%) of polysaccharide from *Ganoderma lucidum* was 6.27% using hot-water extraction. In figure 2, three fractions (F1, F2, and F3) were found by ion exchange chromatography (DEAE-52) and GLP was purified on Sephadex S-300 column.

**Figure 2.**
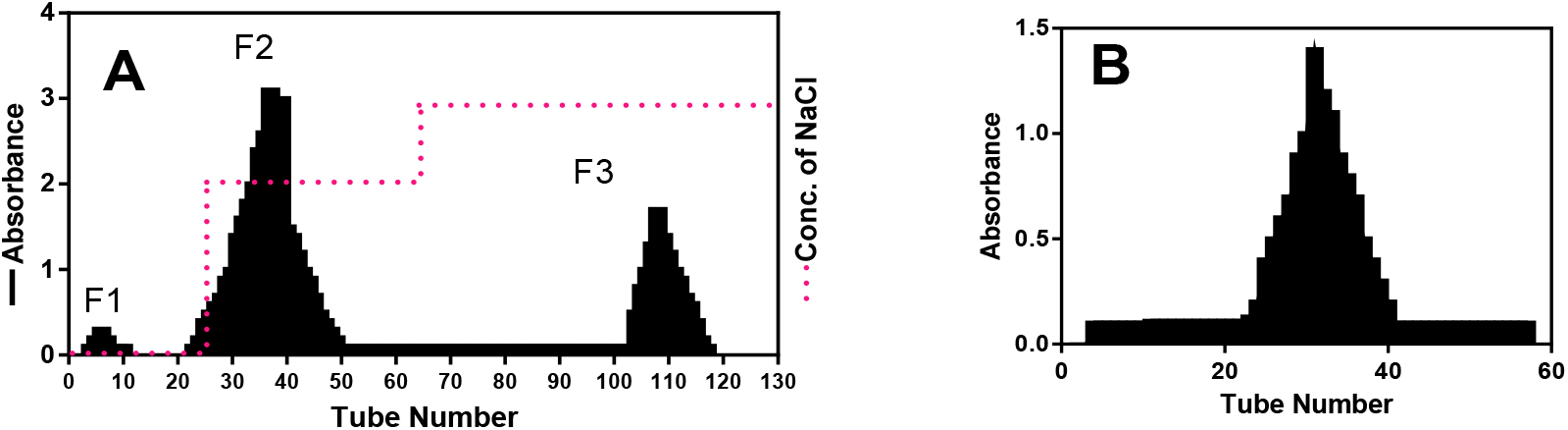
Elution profile of GLP (A) DEAE-52 column and (B) Sephadex S-300 column. GLP; *Ganoderma lucidum* polysaccharide

The molecular weight of GLP was determined using Kav values of GLP and the molecular weight of GLP was 159498.8 (Da) using standard curve (Fig. 3).

**Figure 3.**
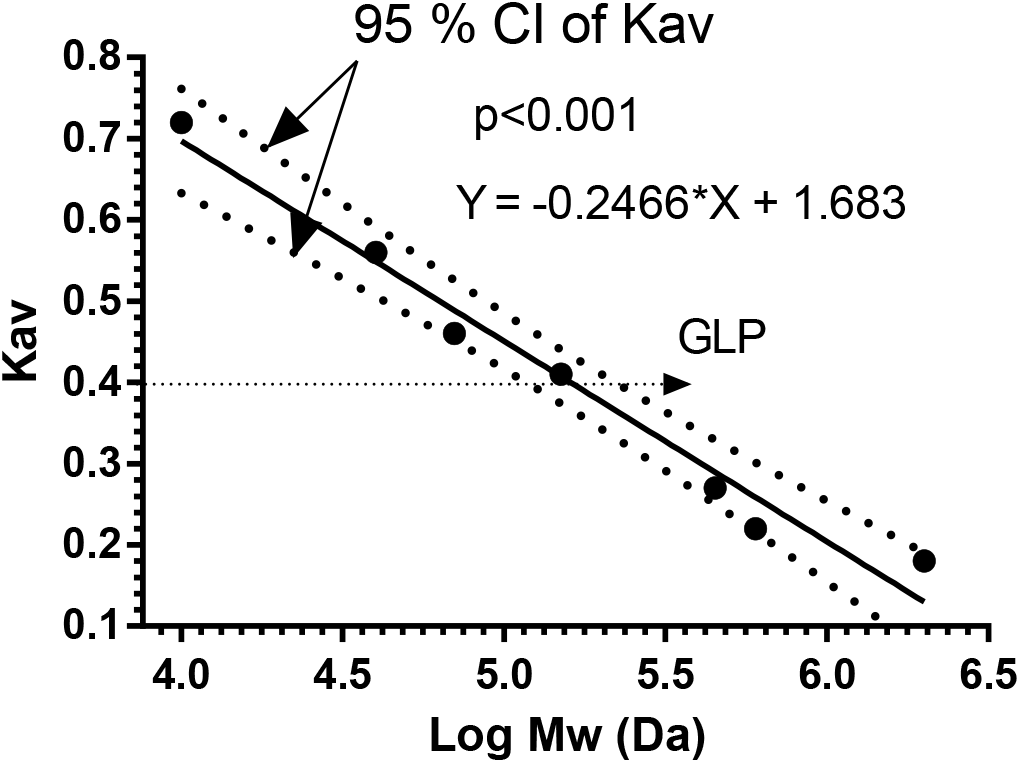
Standard curve of dextran series for the determination of Mw of the polysaccharide of *Ganoderma lucidum* based on Kav.

#### UV-Viz and FTIR analysis

In UV-Viz analysis, the crude GLP fraction had protein (280 nm) impurity (Fig. 4A) and after deproteination, there was no absorbance at 280 nm/ 260 nm. Fig. 4B indicates that the GLP was free from protein and nucleic acids.

**Figure 4.**
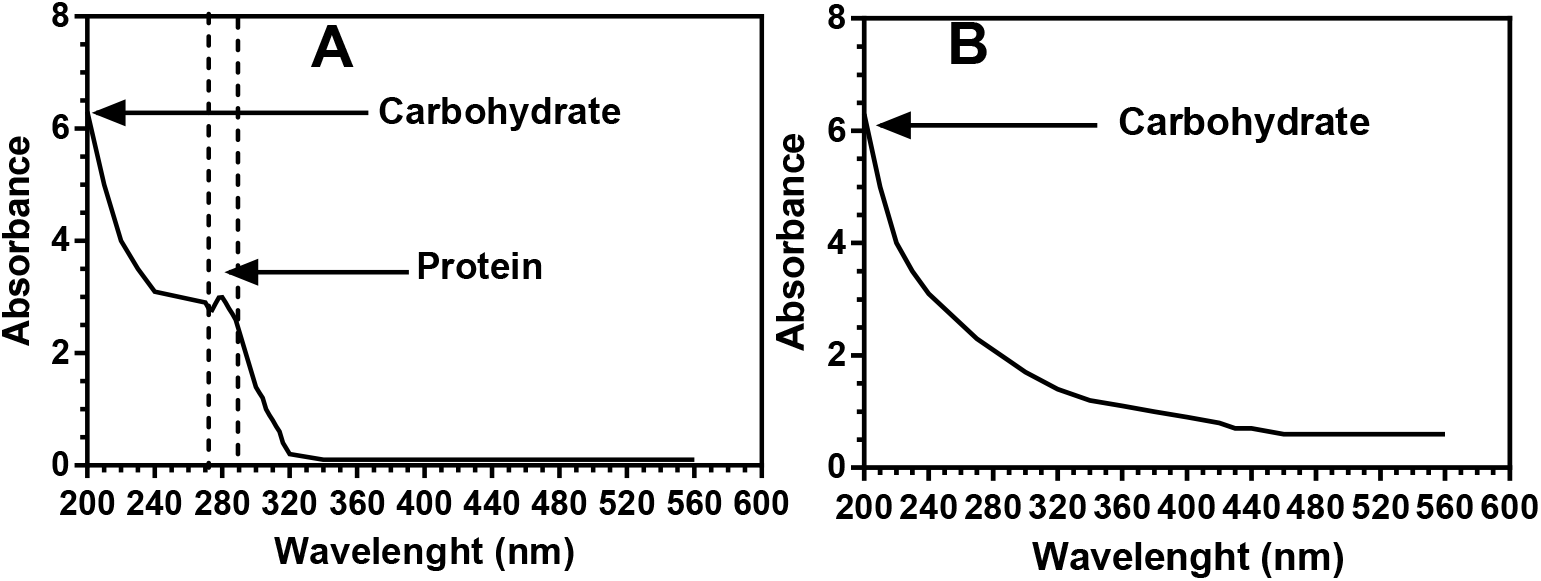
UV-Viz analysis of GLP. GLP; *Ganoderma lucidum* polysaccharide

FT-IR spectroscopy was used to identify characteristic functional groups of purified GLP. FT-IR spectra were reported in the range of 4,000-400 cm^−1^. In figure 5 (C), the FT-IR of GLP showed the characteristic absorbance at about 3419.8 cm^−1^, 2924.5 cm^−1^, and 1615.1 cm^−1^ for −OH groups, C-H bond, and C=O bond respectively.The spectra at 990-1050 cm^−1^ confirmed the presence of β-glycosidic linkage. The main chain of polysaccharide is α-D-glucopyranose.

**Figure 5.**
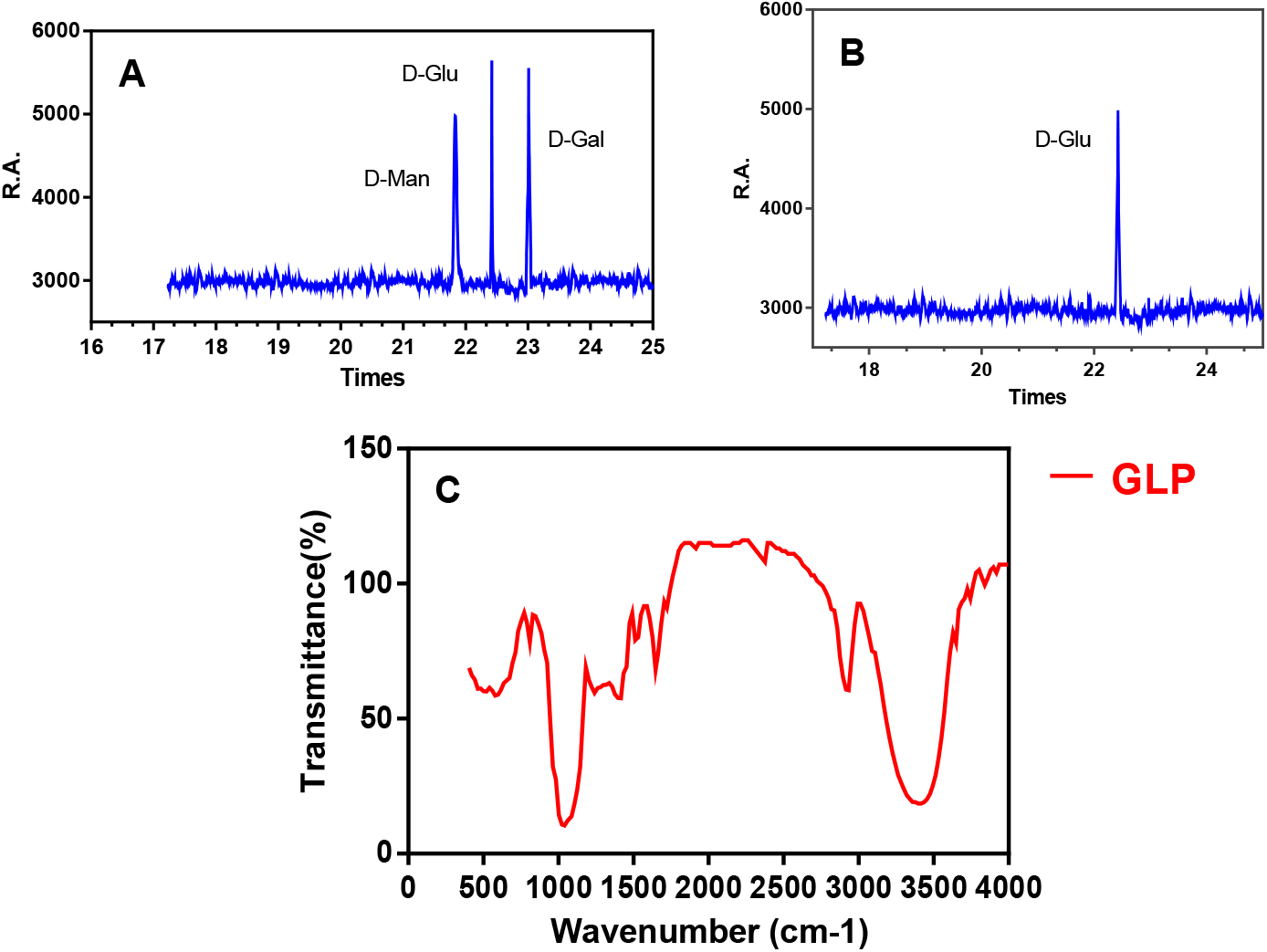
Monosaccharide compositions and FT-IR analysis of GLP. (A) GC results of standard momosaccharides, (B) GC spectra of GLP fraction, and (C) FT-IR spectra of GLP. D-Man; D-Mannose, D-Glu; D-Glucose; and D-Gal; D-Galactose. GLP; *Ganoderma lucidum*

### Monosaccharides compositions

GLP was homopolysaccharides composing of D-glucose. Figure 5<u>5</u> (A, B) shows the GC results of standard momosaccharides and GLP.

### Antioxidant activity screening

Antioxidant activity was performed with the DPPH free radicals scavenging assay. Figure 6(A) and 6(B) illustrate % antioxidant activity of GLP. The IC_50_ alue of GLP was 202.4 μg/mL. It was concentration dependent activity. The highest concentration of GLP showed the maxium<u>maximum</u> effect of free radical scavenging activity against DPPH radicals. There was no significant difference between GLP and Vit-C (>0.05).

**Figure 6.**
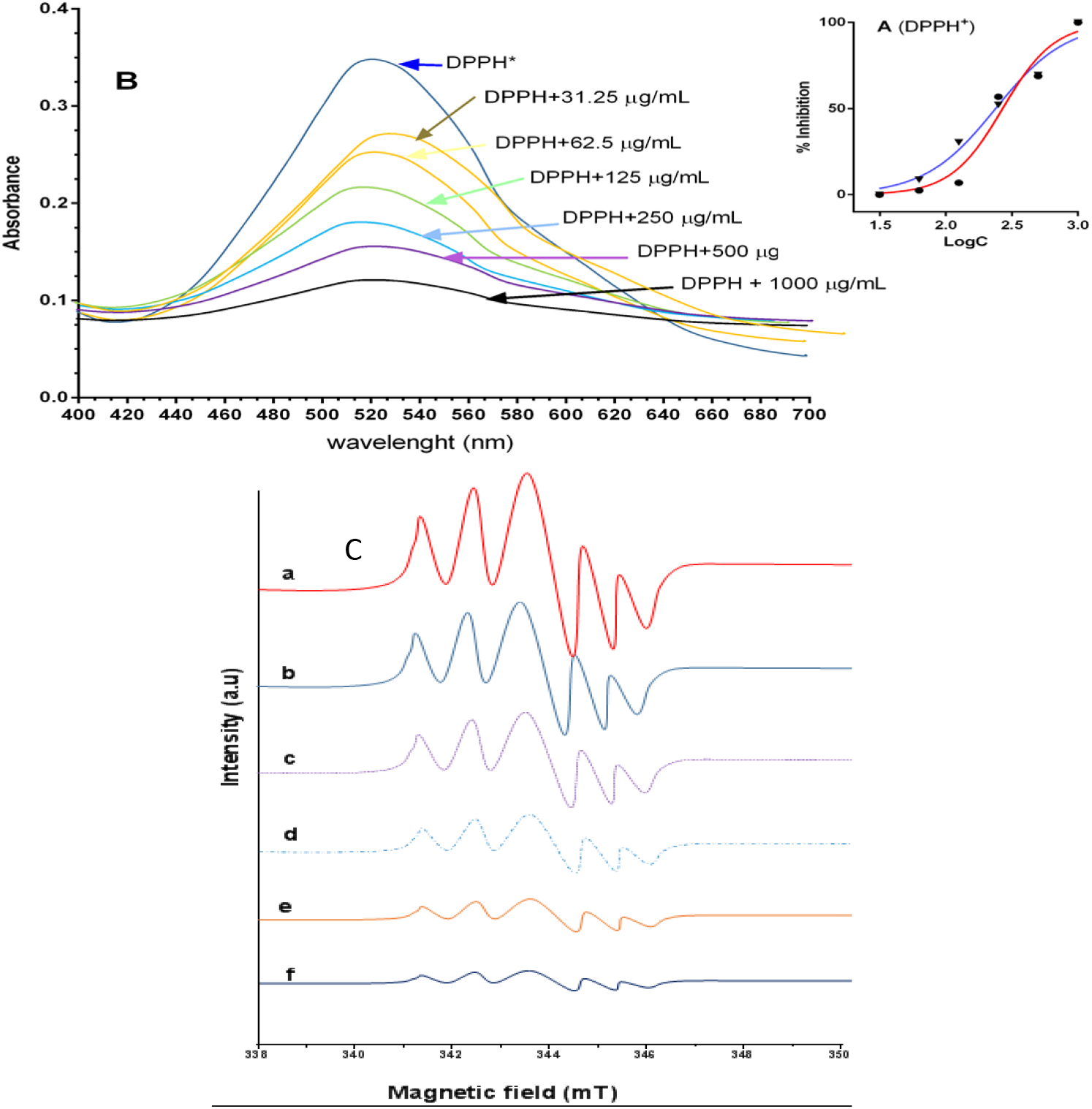
(A) Dose-response curve (B) UV-Viz absorbance intensity of treated and untreated DPPH solutions (C) EPR of the treaated and untreated DPPH solutions.

In the ERP analysis, the intensity of signal (A.U) decreased with increasing concentration of GLP. So, low concentration of GLP reduces DPPH radicals slowly.

### MTT assay

Figure 7 shows the cytotoxic activity of GLP and doxorubin on MCF-7 cells in dose- and time-dependent manner. We examined different concentration of GLP for 48 h and the effective doses were calculated from dose-response curve. The GLP and DR showed significant activity against MCF-7 cell line with an IC_50_ 110.907 μg/mL and 58.206 μg/mL respectively.

**Figure 7.**
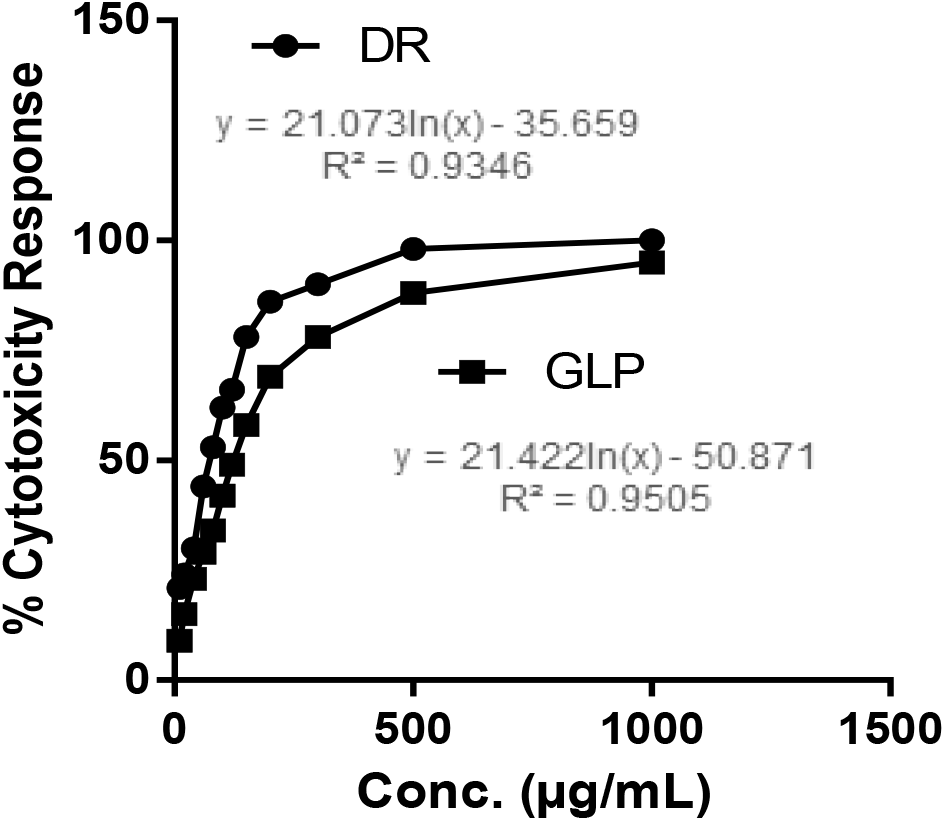
Inhibition of MCF-7 cell proliferation by GLP and DR. *GLP-Ganoderma lucidum* polysaccharide, DR-Doxorubin

### Induction of Apoptosis by cell cycle arrest

The effect of GLP on apoptosis induction in MCF-7 cells was determined using flow cytometry. In this experiments, MCF-7 cells were treated with GLP for 48 h and then prepared to analyze cell cycle phases. The figure 8 illustrated the viable cells, early-phase apoptotic cells, dead cells and necrotic cells of MCF-7. The early apoptotic cells and dead cells of MCF-7 at 2<u>1</u>xIC_50_ were 18.23% and 8.45% respectively (Figure 8C and 8D). At 2xIC_50_, GLP increased the cells in the lower right quadrant and upper right quadrant of MCF-7 to 34.76% and 16.34% respectively (Fig.8A-D).

**Figure 8.**
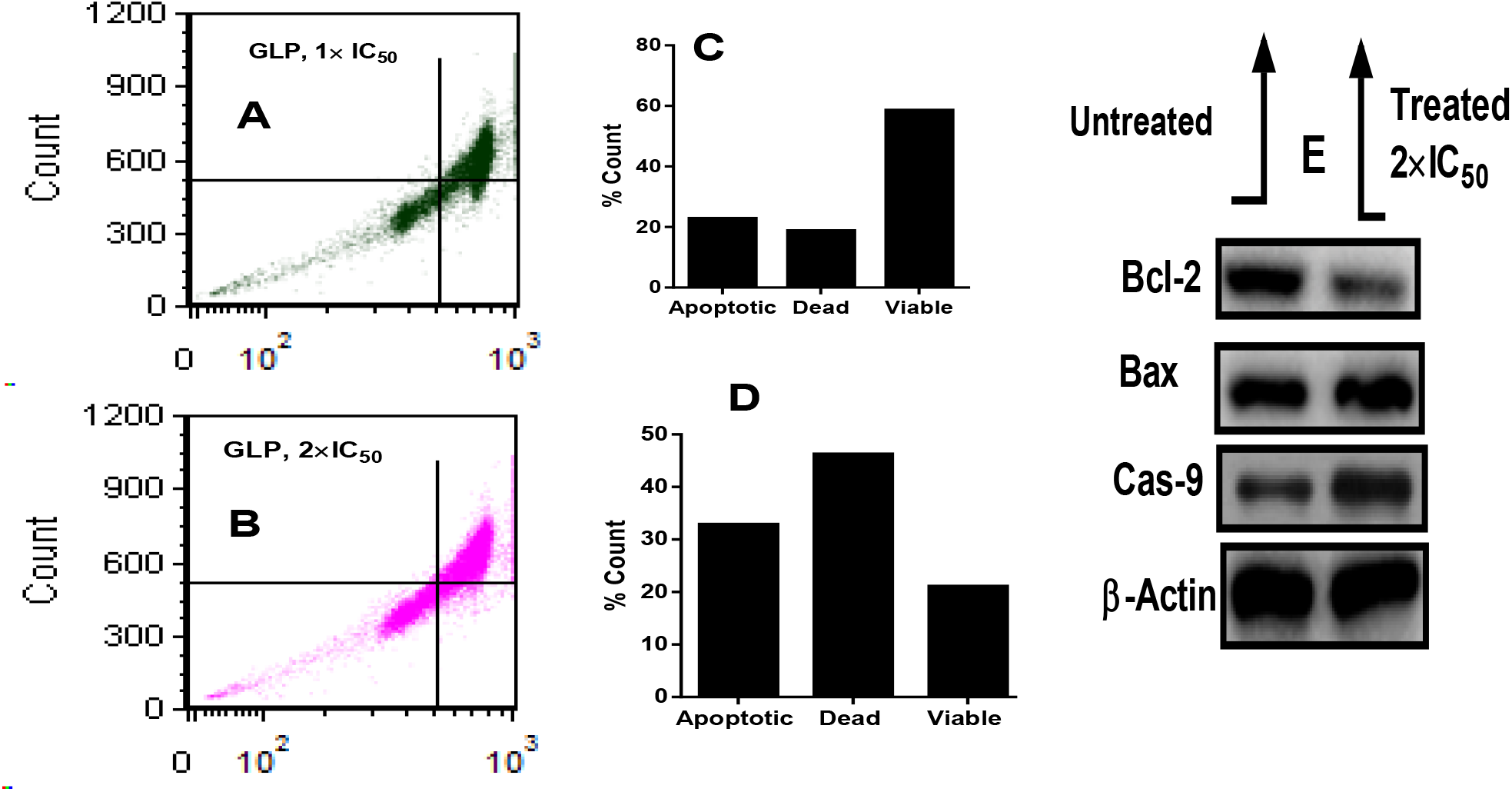
Cell cycle phases distribution of MCF-7 cells using flow cytometry analysis. (A) GLP, 1 x IC_50_ (B) GLP, 2 x IC_50_ (C) Cell count for 1 x IC_50_ GLP (D) Cell count for 2 x IC_50_ GLP and (E) Apoptotic protein expression of Bcl-2, Bax, and Caspase-9 induced by GLP at 2 x IC_50_ on MCF-7 cells.

### Induction of Apoptosis by protein expression

According to figure 8E, the Bcl-2 was down-regulated while Bax gene was up-regulated in treated MCF-7 cells. In addition, GLP induced to increase the expression of Caspase-9. So, GLP significantly increased the expression of Bax and Caspase-9 and suppressed the Bcl-2 in treated MCF-7 cells compared to untreated MCF-7 cells.

## Discussion

The present study shows that the polysaccharide extracted from the fruiting bodies of *Ganoderma lucidum*(GL), termed GLP is homopolysaccharide. The purification of GLP was carried out using DEAE-52 cellulose and Sepharose CL-6B and The purifed polysaccharide was confirmed using UV-Viz and FTIR spectral analysis. GC spectra showed that GLP was a homopolysaccharide. GLP has been shown to consist of D-glucose (Fig.5A and B). The antioxidant homopolysaccharide was confirmed through the DPPH free radical scavenging activities(Fig. 6A and B). Free radicals are derived from normal metabolic processes in the human body and external sources such as industrial chemicals, air pollutants, X-rays, and smoking(20). These are highly reactive species that are capable of damaging biologically significant molecules, such as DNA, proteins, carbohydrates, and lipids(21). Antioxidants can donate an electron to free radicals and neutralize it, resulting delay or inhibit cellular damage(22). Numerous studies have shown that polysaccharides are active mushroom components, which show various pharmacological activities such as antitumor, antioxidant functions(15). In our study, we analyzed and compared DPPH free radicals scavenging activity with that of vitamin C. In DPPH free radical scavenging, it was demonstrated that the IC_50_ value of GLP was 202.4 μg/ml. Furthermore, the antioxidant GLP was used to evaluate the anticancer role in MCF-7 cells line. GLP underwent series of experiments to test their anti-cancer efficacy. Our study was conducted on Breast cancer cells lines. Breast cancer is the most common cancer among women in the world, accounting for 25.4 per cent of the total number of new cases diagnosed in 2018(23). Among breast cancer patients, the average survival of five years was improved by the current treatment methods for patients with breast cancer, but the long-term survival is still poor due to cancer progression and metastasis(24). Breast-conserving surgery, radiation therapy, and mastectomy is the most common treatment for women with early stage breast cancer. Phase III patients undergo chemotherapy with mastectomy, while phase IV patients most often receive radiation, chemotherapy and hormonal therapy(24). There are various subtypes of breast cancer; thus different options are selected for the treatment of breast cancer. The use of chemotherapy drugs is usually associated with the production of deleterious side effects and drug resistance for a longer time(25).

Polysaccharides can be useful in this regard because they are non-toxic and commonly found in natural plants(26). Polysaccharides have reported anti-cancer effects from various sources(10). In our study, GLP showed dose- and time-dependent cytotoxic properties against MCF-7 cells (Figure 7). However, the MCF-7 cells were more sensitive to DR (IC_50_;58.206 μg/mL) than the GLP (IC_50_;110.907 μg/mL). The cytotoxic role of GLP is dose- and time-dependent. Inhibition of cancer progression often involves the regulation of signal transduction resulting in cell growth arrest and apoptosis. Polysaccharides have been studied in the cell lines of MCF-7 in relation to the protein expression of Bax, Bcl-2 and Caspase-9 in apoptosis (Figure 8E). In this study, GLP showed the upregulation of Bax and Caspase-9 protein expression in MCF-7. Apoptosis characterized for the removal of damaged cells or tumor cells by distinct biochemical factors(27). Apoptosis is an extremely controlled and well maintained cell death cycle during which a cell is subjected to self-destruction(28). In multicellular organisms, it is an important mechanism that removes unnecessary or superfluous cells during growth, or neutralizes potentially harmful cells(29). Apoptosis regulation is essential for the maintenance of natural cellular homeostasis. Nevertheless, apoptosis deregulation has been related to various pathologies such as chronic inflammation, cancer and neurodegenerative diseases(30). Apoptosis is a well defined mechanism that displays distinctive morphological and biochemical characteristics(31). They are distinguished by cell shrinking, membrane blebbing, chromatin condensation and nuclear fragmentation, and by the development of apoptotic bodies digested by macrophages. Apoptotic pathways lead to cysteine dependent aspartate proteases (caspases) being activated(32). Caspase-9 is the initiator caspase linked to the apoptosis intrinsic or mitochondrial pathway(33). When triggered, caspase-9 cleaves and triggers downstream effector caspases 3 and 7, targeting important regulatory and structural proteins for proteolysis to effect cell death(34). Lee *et al*. reported that the expression level of anti-apoptotic proteins (Bcl-2) may be decreased by plant phytochemicals(35). The Bax protein triggers the release of cytochrome C and other pro-apoptotic proteins into cytosol from the outer mitochondrial membrane. Bax further triggers apoptosis initiator Caspase-9 by the Caspase-3 cleavage resulting in apoptosis induction(36). From our results, GLP induce apoptosis through the inhibition of Bcl-2 protein expression and upstream regulation of Bax and Caspase-9 expression (Figure 8E).

Previous studies have identified a comprehensive relationship between different fractions of polysaccharides from *Ganoderma lucidum (10)*, *Lentinus edodes*, *Lycium barbarum(37)*, *Ganoderma lucidum* and *Borojoa sorbilis* cuter(4) in different models of animals. Our results have concluded that mushroom polysaccharide, as a potential anti-cancer agent for various anti-cancer treatments, could be employed as the main compound of natural plant products based on the promotion of cell cycle arrest, apoptosis and inhibition of MCF-7 proliferation in *vitro*. However, earlier studies have identified a comprehensive relationship between different fractions of polysaccharides in various cancer(38). In *vivo* trials using tested polysaccharide GLP at the next stage will deliver their detailed efficacy study for better anti-cancer prediction.

## Conclusion

Our research aimed to evaluate the apoptosis induction in MCF-7 cells by GLP derived from the *Ganoderma lucidum*. Because apoptosis has a critical role in the homeostasis of tissues and the prevention of cancer growth, it is now an important goal for cancer treating and preventing the regulation of cell production and the activation of cell apoptosis. GLP was extracted with hot water and further, separation and purification of GLP was carried out using DEAE-52 cellulose and Sepharose CL-6B respectively. GLP was a homo-polysaccharide consisting of D-glucose without protein. According to the degree of purity of GLP was tested and detected by UV-viz and FTIR spectral analysis. GLP had good antioxidant activity with the IC_50_ values of 202.4 μg/mL in DPPH. MTT assay measured the cytotoxicity of GLP, and results showed that GLP significantly inhibited the growth of MCF-7 cells. Such results indicated the GLP could be investigated as a possible anti-tumor agent. The combined data suggest that GLP inhibits the growth of MCF-7 breast cancer cells by inducing the cell cycle arrest and apoptosis. In this study, GLP-induced apoptosis in breast cancer cells demonstrated a dose- and time-dependent manner of activity. GLP induce apoptosis by the down-regulation of Bcl-2 and the up-regulation of Bax and Caspase-9. We assume GLP has the potential anticancer role; more research is needed in animal tumor models to confirm *in vivo* anti-cancer activity of GLP.

